# Disrupted excitation-inhibition balance in cognitively normal individuals at risk of Alzheimer’s disease

**DOI:** 10.1101/2023.08.21.554061

**Authors:** Igor Fortel, Liang Zhan, Olusola Ajilore, Yichao Wu, Scott Mackin, Alex Leow

## Abstract

**Background:** Sex differences impact Alzheimer’s disease (AD) neuropathology, but cell-to-network level dysfunctions in the prodromal phase are unclear. Alterations in hippocampal excitation-inhibition balance (EIB) have recently been linked to early AD pathology.

**Objective:** Examine how AD risk factors (age, APOE-ɛ4, amyloid-β) relate to hippocampal EIB in cognitively normal males and females using connectome-level measures.

**Methods:** Individuals from the OASIS-3 cohort (age 42-95) were studied (N = 437), with a subset aged 65+ undergoing neuropsychological testing (N = 231).

**Results:** In absence of AD risk factors (APOE-ɛ4/Aβ+), whole-brain EIB decreases with age more significantly in males than females (p = 0.021, β = -0.007). Regression modeling including APOE-ɛ4 allele carriers (Aβ-) yielded a significant positive AGE-by-APOE interaction in the right hippocampus for females only (p = 0.013, β = 0.014), persisting with inclusion of Aβ+ individuals (p = 0.012, β = 0.014). Partial correlation analyses of neuropsychological testing showed significant associations with EIB in females: positive correlations between right hippocampal EIB with categorical fluency and whole-brain EIB with the trail-making test (p < 0.05).

**Conclusion:** Sex differences in EIB emerge during normal aging and progresses differently with AD risk. Results suggest APOE-ɛ4 disrupts hippocampal balance more than amyloid in females. Increased excitation correlates positively with neuropsychological performance in the female group, suggesting a duality in terms of potential beneficial effects prior to cognitive impairment. This underscores the translational relevance of APOE-ɛ4 related hyperexcitation in females, potentially informing therapeutic targets or early interventions to mitigate AD progression in this vulnerable population.

## Introduction

With networks composed of billions of neurons and synaptic connections, the human brain is a highly complex dynamical system wherein neural populations coordinate electrical activity to balance excitatory and inhibitory neural activity across different brain regions. Balanced excitation and inhibition is crucial for adjusting neural input/output relationships in cortical networks, regulating their dynamic range of responses to stimuli [1], and achieving optimal information capacity and transfer [2]. Moreover, maintaining this balance is essential for healthy cerebral activity, homeostatic control, and potentially contributing to lifespan longevity [3–5]. Imbalances, however, may lead to serious neurological disorders such as autism, characterized by a shift towards a more excitatory state [6,7], and Down syndrome, linked to increased inhibition (or decreased excitation) [8].

Growing evidence supports the hypothesis that neuronal hyperexcitation, due to the susceptibility of inhibitory GABAergic interneurons to Apolipoprotein (APOE) ε4-mediated neurotoxicity (particularly in the hippocampal dentate gyrus), may represent the earliest changes indicative of Alzheimer’s disease (AD) pathology [9–11]. Normal age-related declines in excitatory connectivity may result from alterations in synaptic plasticity, receptor expression, and neuronal morphology, leading to changes in excitatory and inhibitory signaling [9–11]. Here, we leverage a recently developed framework based on a pairwise maximum entropy model from statistical physics to computationally infer excitation-inhibition balance (EIB) from combined structural and functional MRI imaging data [12]. Inspired by the Ising spin-glass model representation of brain dynamics, self-organized patterns of connectivity form through the spontaneous fluctuations of random spins [13]. Spin-glass models have been used to characterize both complex microscale dynamics of the human brain [14–17] and macroscale interactions [18–23]. In fact, unconstrained maximum entropy models have been shown to accurately represent spatiotemporal coactivations in neuronal spike trains [23–26] and patterns of BOLD activity [26–29] . Recently, Zanoci and colleagues [30] demonstrated that the Ising model can capture collective neuronal behavior during wakefulness, light sleep, and deep sleep when both excitatory (E) and inhibitory (I) neurons are modeled.

Sex differences play an important role in AD risk, with females being more susceptible to APOE-related pathogenic changes than males [31,32]. Among other genetic and epigenetic factors, this risk may be attributed to hormonal differences, such as the protective effects of estrogen on neuronal function and synaptic plasticity, which decline during menopause in females, potentially increasing their susceptibility to APOE-related pathogenic changes like hyperexcitation [33,34]. However, the field of clinical AD often treats sex as a covariate, limiting the potential for discovering interacting effects regarding disease progression [35]. Recent studies from the Alzheimer’s Disease Neuroimaging Initiative (ADNI) suggest that approximately seventy-five percent of patients with AD are female and appear to progress to AD more often than males [36]. While the subjects of this study are unimpaired, our hypothesis is that our metric describing EIB can detect subtle shifts in hippocampal dynamics related to APOE or amyloid-β influences.

We apply our computational framework on a large cohort (OASIS-3: Open Access Series of Imaging Studies [37]) of individuals aged 42 to 95 years without cognitive impairment. The first aim is to test the hypothesis that female ε4 carriers would exhibit connectome signatures of hyperexcitation within hippocampal regions compared to non-ε4 carriers, which may confer a greater vulnerability to Alzheimer’s disease neuropathology. Moreover, we hypothesize that even females without elevated risk of developing AD (either through APOE4 or Aβ+) also have diverging EIB compared to men, contributing to higher risk of APOE or amyloid-mediated effects of hyperexcitation. Lastly, we investigate sex differences in EIB concerning neuropsychological testing with a focus on hippocampal dynamics, which play distinct roles in detecting and monitoring cognitive and functional changes associated with dementia-associated diseases [38].

While studies assessing the effects of sex and APOE4 or Aβ+ on specific MRI-based BOLD signal changes have produced varied results, early identification and intervention are critical to improving patient outcomes. We aim to expand on these sex differences, providing new insights and suggesting that a network with spatiotemporal structure may help in better understanding AD progression and identify subtle changes to neural dynamics before the onset of irreversible degeneration. This can lead to more informed therapeutic targets and early interventions to mitigate AD progression in vulnerable populations, potentially guiding the development of tailored, effective treatments and improving early diagnosis. Enhanced understanding of EIB dynamics could contribute to personalized medicine approaches in AD, ultimately leading to better patient outcomes.

## Materials and Methods

### Participants

The Open Access Series of Imaging Studies (OASIS-3) is a retrospective compilation of over 1000+ participants with imaging data collected over the course of 30 years across several ongoing projects at the Charles F. and Joanne Knight Alzheimer Disease Research Center at Washington University in St. Louis. These projects include the Memory and Aging Project, Adult Children Study, and Healthy Aging and Senile Dementia. In total, OASIS-3 comprises a diverse collection of clinical, neuropsychological, neuroimaging, and biomarker data for 1098 participants (age: 42–95 years). It is important to note that not all participants completed the full protocol, and only individuals aged 65 or older completed neuropsychological testing [39].

Thus, in this study we analyzed a cross-section of individuals with combined structural MRI, functional MRI, and PET imaging sessions (N = 437), including apolipoprotein E gene (APOE) genotyping, extracted from (https://central.xnat.org). In this study, genotype was analyzed as a dichotomous variable defined as APOE4 non-carriers (ε3/ε3 allele combination) and APOE4 carriers (ε3/ε4 or ε4/ε4 allele combinations) since the ε4 allele is commonly associated with an increased AD risk [40–42]. Herein, NC will refer to APOE4 non-carriers, and APOE4 will refer to carriers of at least one ε4 carrier.

For the purposes of this study, we included only participants classified as cognitively normal (no impairment) according to the Clinical Dementia Rating (CDR) scale (CDR = 0). Inclusion of only cognitively normal participants (CDR = 0) ensures that the analysis focuses on individuals without any impairment, providing a clearer picture of the factors affecting healthy aging. We further excluded participants carrying one or more APOE-ε2 alleles due to the neuroprotective effect associated with this allele against AD-related neurodegeneration [43,44]. Consequently, this group could not be considered as part of either the control or carrier groups. Exclusion of participants carrying one or more APOE-ε2 alleles eliminates potential bias introduced by the neuroprotective effect of this allele, resulting in a more accurate assessment of the impact of other factors on AD risk [45,46].

We matched each MRI-PET session with the closest clinical session. For each participant, the first MRI-clinical-session pair with an absolute time difference of < 1 year was selected as the baseline session. In cases where duplicate data were available for subjects, we used the earliest or initial MR imaging session. Using the earliest or initial MR imaging session for participants with duplicate data reduces the potential for confounding effects of disease progression or other time-dependent factors on the analysis. The aggregated participant information used in this study is presented in Table 1.

**Table 1.**
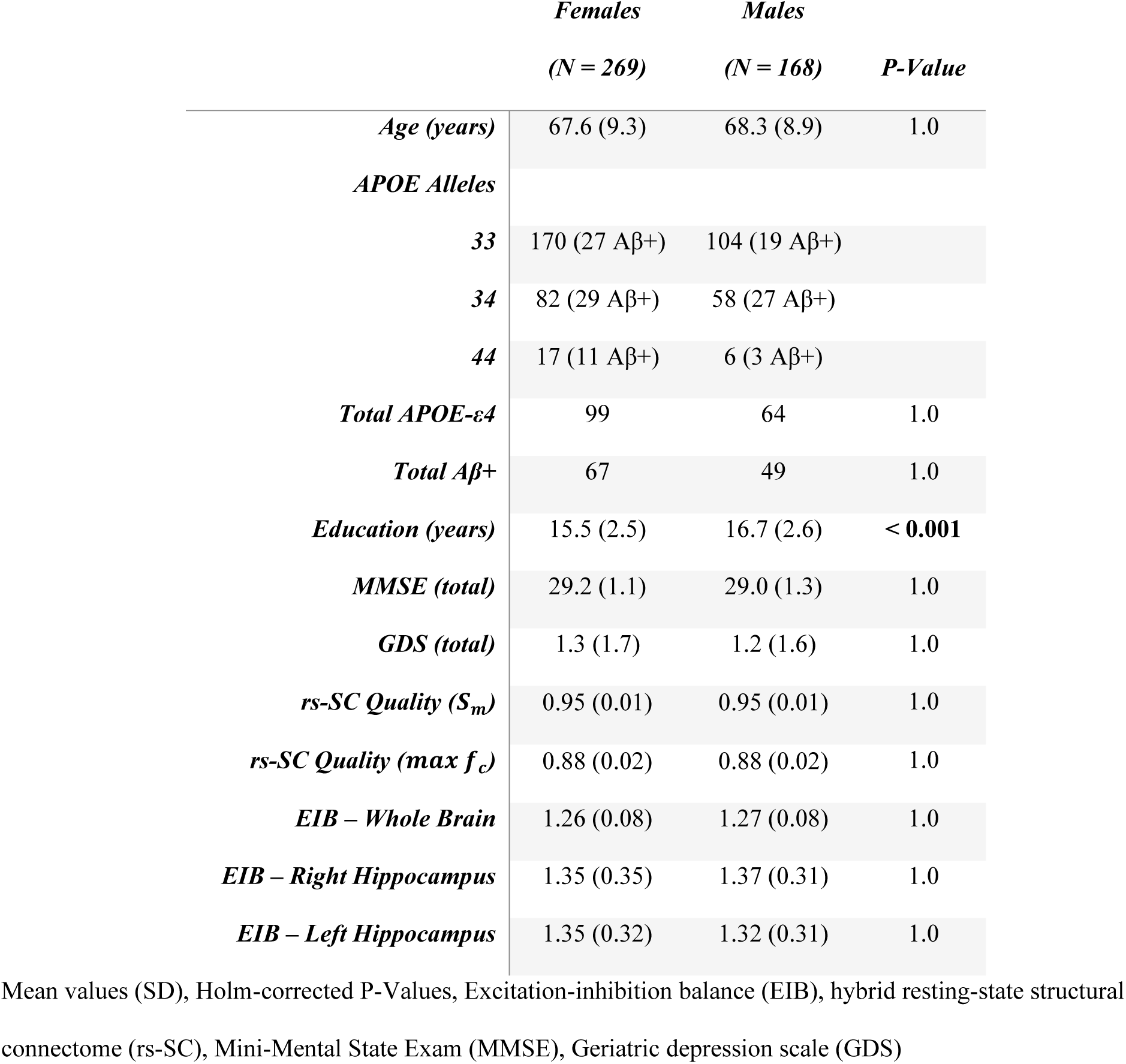
Demographics and screening measures of participants from OASIS-3 [39] used in this study. All subjects are cognitively normal with CDR = 0.

### Cognitive assessment measures and neuropsychological testing

In this study, three screening measures were included to ensure that included subjects are without cognitively impairment: the Mini-Mental Status Examination (MMSE), Geriatric Depression Scale (GDS) and the Clinical Dementia Rating (CDR). The MMSE is a screening tool used to assess global mental status. It is comprised of an 11-question measure (scored from 0 to 30) assessing orientation, attention and concentration, registration, recall, and language. Scores below 24 have been shown to be effective in identifying individuals with possible cognitive impairment [47]. The GDS is a 30-item measure that assesses symptoms of depression in middle-aged and older populations. Scores above 10 indicate possible Major Depressive Disorder [48]. The standardized collection of OASIS-3 was developed at the Alzheimer’s Disease Centers (ADC) program of the National Institute on Aging (NIA) as a central component of the Uniform Data Set (UDS) [49,50]. Here, the clinical labeling of dementia severity was determined based on the Clinical Dementia Rating (CDR). The CDR is a semi-structured interview intended to provide a global dementia severity rating, which is commonly employed for staging and tracking decline in AD [51,52]. The score here represents multiple levels of functional impairment (0 = no impairment; 0.5 = questionable or very mild impairment; 1 = mild impairment; 2 = moderate impairment) and summarizes the estimate of dementia severity [53,54]. As mentioned in the previous section, this study comprises only of individuals with a CDR equal to zero (signifying no impairment).

Further, the protocol used in the OAISIS-3 for neuropsychological testing consists of a battery based on the Uniform Data Set (UDSNB), which was developed in Alzheimer’s Disease Centers (ADC) to establish a standardized data collection system [55]. Participants aged 65 and older completed a battery of neuropsychological tests at the ADRC to evaluate several tests measuring attention, working memory, executive function, processing speed, language and episodic memory. Logical Memory - Story A, a subtest of the Wechsler Memory Scale-Revised measures episodic memory. Participants recall as many details as they can from a short story containing 25 bits of information after it is read aloud by the examiner (LM-I) and again after a 30-minute delay (LM-II, with scores ranging from 0 (no recall) to 25 (complete recall). Semantic memory and language were measured using the Category Fluency test, requiring participants to name as many words belonging to a category, animal and vegetable, in 60 seconds, and the Boston Naming Test, in which participants name drawings of common objects. Psychomotor speed was measured using the WAIS-R Digit Symbol test and the Trail Making Test Part A. The WAIS-R Digit Symbol test is scored on the number of digit symbol pairs completed in 90 seconds. Executive function was measured using the Trail Making Test Part B. In the Trail Making Test participants were asked to make a trail by connecting a series of numbers (1-26) for part A and connecting a series of alternating numbers and letters (1-A-2-B) for part B. The outcome measure used in this manuscript is the total time to complete in seconds with a max of 150s for Trails A and 300s for Trails B. A complete description of cognitive assessment measure and neuropsychological testing completed for the OASIS-3 dataset can be found in the data dictionary (https://www.oasis-brains.org/)

### MRI and PET imaging

MRI and PET imaging used in this study from the OASIS-3: Longitudinal Neuroimaging, Clinical, and Cognitive dataset for Normal Aging and Alzheimer’s disease [37] was collected in a 16-channel head coil of a Siemens TIM Trio 3T scanner. In this study, only cognitively normal individuals with structural (Diffusion-weighted imaging and T1-weighted imaging), resting-state functional (fMRI) and PET imaging were included. Processing of the MRI imaging is described in the supplementary materials detailing the processing of structural and functional networks, as well as the procedure for constructing a hybrid resting-state structural connectome. In short, our team recently combined MRI-derived multimodal connectome data (resting-state functional and structural) with statistical physics to obtain a macroscopic account of inhibitory and excitatory long-range connections [12,56]. Here, we employed the Ising model, a well-known spin-glass model from statistical physics in which the states, also referred to as “spin configurations”, of interacting units – in our case brain regions connected by white matter edges – are constrained to be either 1 (“active”) or -1 (“inactive”). As described previously [12,56], we construct a function-by-structure embedding (FSE) using a constrained pseudolikelihood estimation technique wherein pairwise interaction coefficients are inferred from the observed data (BOLD time series) and underlying structural connectome, resulting in a spatiotemporal network describing both structure and function (called the hybrid resting-state structural connectome, or rs-SC).

Note that although our framework for constructing the rs-SC does not explicitly model brain activity at a neuronal level, several seminal studies have used the Ising model to investigate neuronal firing patterns with success, showing that ensemble level activities can be well explained by pairwise Ising interactions [24,57]. To be clear, from a connectomics perspective, if several brain regions are identified to have an increase in positive edges in the rs-SC, collectively, that would suggest a wider-spread pattern of coupling (i.e., more likely to exhibit a pattern of global coupling) that may subserve hyperexcitation. It is in this context that we conceptualize the Excitation-Inhibition (E-I) ratio, a global (whole-brain) or local (ROI-level) estimation of EIB, computed as the sum of positive edges divided by the sum of negative edges. For example, if a brain region in the network has 45 positive interactions (edges) and 34 negative interactions with other brain regions, then the E/I ratio = 45/34, or 1.32 (a value of 1 indicates perfect EIB).

PET imaging included both [11C]-Pittsburgh compound B, as well as Florbetapir ([18F]-AV45). The PET image processing steps can be found in the data dictionary (https://www.oasis-brains.org/). For this study, we utilized the partial volume corrected mean cortical Standard Uptake Value Ratio (SUVR) with regional spread function (SUVR rsf) provided in the dataset database (https://central.xnat.org). To assess global amyloid burden based on the PET imaging, the mean cortical SVUR was defined as the arithmetic mean of SUVRs from the precuneus, prefrontal cortex, gyrus rectus, and lateral temporal regions (include citations). A positive amyloid-β PET scan (Aβ+) was defined as having a mean cortical SUVR rsf > 1.42 for [11C]-Pittsburgh compound B PET or SUVR rsf > 1.19 for Florbetapir PET [39,58]. For simplicity with two different PET tracers utilized in this dataset, in the ensuing analyses we treat Aβ as a dichotomous variable wherein an individual may be either Aβ- or Aβ+.

### Statistical analysis

Demographic, screening and clinical measures for males and females were compared using the Wilcoxon rank-sum tests for continuous variables and Fisher’s exact test for categorical variables. These non-parametric tests were chosen due to their robustness to non-normality and applicability to small or unbalanced sample sizes. The statistical tests were initially used to identify differences in demographic, screening, or clinical measures between males and females. However, the decision to include these variables as covariates in the regression models was also based on their potential impact on the outcome variables as informed by existing literature. This consideration led to the addition to the AD risk factors being evaluated (ε4 carriers (APOE)/non-carriers (NC), sex, age, and amyloid-β positivity (Aβ+/Aβ-).

Whole-brain and hippocampal EIB were evaluated using linear regression models, which allowed us to estimate the relationships between the predictors and the outcome variables, as well as assess potential interaction effects. Three separate regression models were tested (with sex = 1 for females and sex = 0 for males): The first regression model examined EIB as a function of age and sex interactions for cognitively normal males and females without any known predispositions to AD neuropathology (i.e., individuals with CDR = 0, Aβ-, and APOE-ε3/ε3 allele combinations). This model aimed to identify and evaluate the progression of EIB between sexes under normal-aging conditions (N = 228, 143 Female) without any known genetic risk factors for AD. The second regression model assessed EIB as a function of age and APOE interactions for individuals that are Aβ-to evaluate the influence of the APOE4 allele on EIB in healthy aging (N = 321, 202 Female). APOE is a dichotomous variable indicating individuals with at least one APOE4 allele or none (ε3/ε3). The third regression model examined EIB as a function of the interactions between age, APOE, and amyloid (allowing for up to 4-way interactions between predictors), as well as two-way interactions of APOE – amyloid when controlling for other confounds (N = 437, 269 Female). Amyloid positivity is a dichotomous variable (Aβ+/Aβ-) based on thresholding of SUVR data from PET imaging, as described in the methods section. This model was analyzed separately for each sex to investigate the effect of ε4 and amyloid on EIB specific to males and females.

Lastly, we tested for relationships between cognitive measures, neuropsychological tests, and EIB for each sex using partial correlation analysis to control for age, APOE, and amyloid as confounds. Partial correlation analysis was chosen as it allows for the evaluation of the association between two variables while controlling for the effects of other variables. We further performed correlation analysis to test for associations between EIB and screening measures (MMSE, GDS, Education). P-values were adjusted for multiple comparisons using the Holm correction to control the family-wise error rate and reduce the likelihood of false positives. A p-value < 0.05 was considered statistically significant, and p < 0.1 was considered marginally significant. Analyses were conducted in MATLAB R2021b.

## Results

### Computationally inferring excitation-inhibition balance from MRI imaging data

We compute excitation-inhibition balance (EIB) for the whole-brain as well as for left and right hippocampal regions as determined by the Harvard-Oxford Atlas used in the processing of the MRI imaging. Here we follow the three-stage process for obtaining an optimized hybrid resting-state structural connectome (rs-SC) via a function-by-structure (FSE) embedding. To evaluate the quality of the generated rs-SC, we compute the similarity metric and evaluate the quality of reconstructing observed FC. As described in the supplement, the similarity metric S_m_ evaluates the constraint and is computed as the correlation between the magnitude of rs-SC and the structural connectome. The max f_c_ metric measures the quality of reconstructing observed FC using Monte Carlo simulations of the Ising model with our rs-SC being used to define the interaction network. The value for structural similarity (S_m_) in this dataset is 0.9, the max is 0.98, with a mean of 0.95 and variance of 0.0001. The minimum correlation between observed and simulated FC (max f_c_) in this dataset is 0.82, the maximum is 0.93, the mean is 0.88 with variance of 0.0004. This suggests that the FSE framework works well in constructing the hybrid connectome with low inter-subject variability. From the rs-SC, we compute E-I by summing the positive edges in the network (excitatory interactions) divided by the negative edges in the network (inhibitory interactions). This is computed for the whole brain (105 ROIs) as well as isolating the bilateral hemispheres of the hippocampus (left and right) for further analyses.

### Excitation-inhibition balance of cognitively normal individuals without APOE4 or significant amyloid

With EIB computed for each subject at the whole brain and hippocampal levels, we first evaluate sex differences in EIB for cognitively normal individuals without known risks of developing AD (e.g., APOE-ε3/ε3 and Aβ-). A subset of subjects from table 1, we note only a statistically significant difference in male (N = 85) and female (N = 143) education (in years) with Holm-corrected P < 0.001, noting that on average the females in this group have less education (15.5 years) as compared to males (16.7). As a result, we account for education in our regression models due to significant group difference between males and females. In figure 1 we present the progression of global EIB and hippocampal EIB (left and right). Here, linear regression analysis evaluating whole-brain EIB with interaction effects of age x sex result in p = 0.021 (β = -0.002, SE = 0.001), suggesting that males may have a greater increase in global inhibitory tone than females (or decrease in excitatory interactions). No difference is found for this subset of the OASIS-3 cohort in terms of hippocampal EIB. The result presented in the figure suggests that females maintain EIB from middle age to old age better than men, potentially a result of some compensatory mechanisms to a decline in excitatory tone. Regression results including estimates, p-values and standard error for all main and interaction effects are shared in Supplemental Table 1. Sensitivity analysis was performed using Cook’s Distance for outlier detection in our model. Exclusion of this outlier resulted in only marginal changes in our regression analysis without significant changes to results or overall conclusion, suggesting that the influence of this single data point on the estimated regression line is minimal. Therefore, the inclusion or exclusion of this outlier did not significantly impact the overall results, lending further credibility to our conclusion that men may have a greater increase in global inhibitory tone than women (or decrease in excitatory interactions) in the absence of AD risk factors.

**Figure 1.**
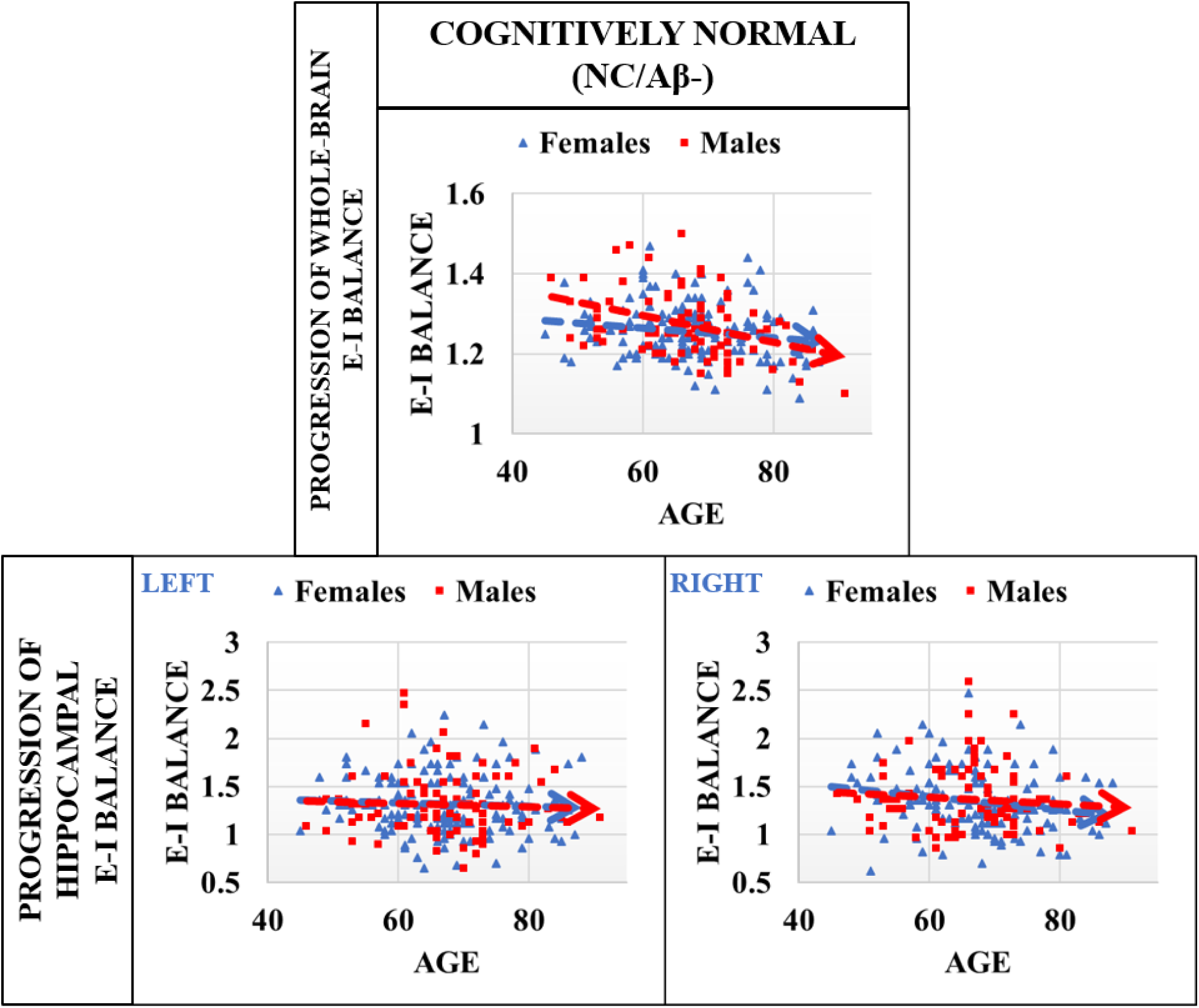
Progression of EIB in cognitively normal males and females without AD risk factors (APOE-ε3/ε3, and Aβ-). Progression of global EIB suggests that males may have a greater increase in inhibitory tone than females with age (or decrease in excitatory interactions). Linear mixed effects regression analysis evaluating the interaction effects of age x sex result in **p = 0.021** (β = -0.002, SE = 0.001). No group differences were identified in the right or left hippocampus. Results suggest that EIB is more stable in females than males with increasing age (in absence of AD risk factors), perhaps due to some underlying compensatory mechanisms.

### Influence of APOE-ε4 on excitation-inhibition balance

Additionally in this study we aimed to evaluate on a larger scale, findings from our previous investigations of sex differences in EIB with respect to the APOE4 allele [12,59], namely that cognitively unimpaired females with at least one APOE4 allele exhibit shifts in excitatory interactions which may confer greater vulnerability to AD neuropathology. Specifically, here we evaluate the progression of EIB with respect to age for whole-brain and hippocampal regions for males and females separately to elucidate the impact of the APOE genotype. Presented in Figure 2 is a comparison of hippocampal EIB progression for cognitively normal males (N = 119) and females (N = 202) that are Aβ-. While our data is not age or sex-matched as in our previous work, figure 2 includes whole-brain comparisons for male and female APOE carriers and non-carriers (NC) based on linear regression analysis. While there is a group difference in whole-brain EIB between female APOE carriers and non-carriers, the difference can be partially explained by age differences in this cross-section, resulting in a p-value of p = 0.061 (β = 0.02, SE = 0.01) for the main effect of APOE status. Visually, only the right hippocampus diverges for female APOE4 carriers as compared to non-carriers. Linear regression analysis evaluating the interaction effects of age x APOE for females results in p = 0.013 (β = 0.014, SE = 0.005). No significant group difference was identified in whole-brain dynamics for either sex, the left hippocampus for women, or for either side of the hippocampus for men. This result suggests that cognitively normal females that are APOE4 carriers and amyloid negative exhibit an increasing effect with age in the EIB of the right hippocampus as compared to non-carriers in the absence of significant amyloid. Regression results including estimates, p-values, and standard error for all main and interaction effects are shared in Supplemental Table 2.

**Figure 2.**
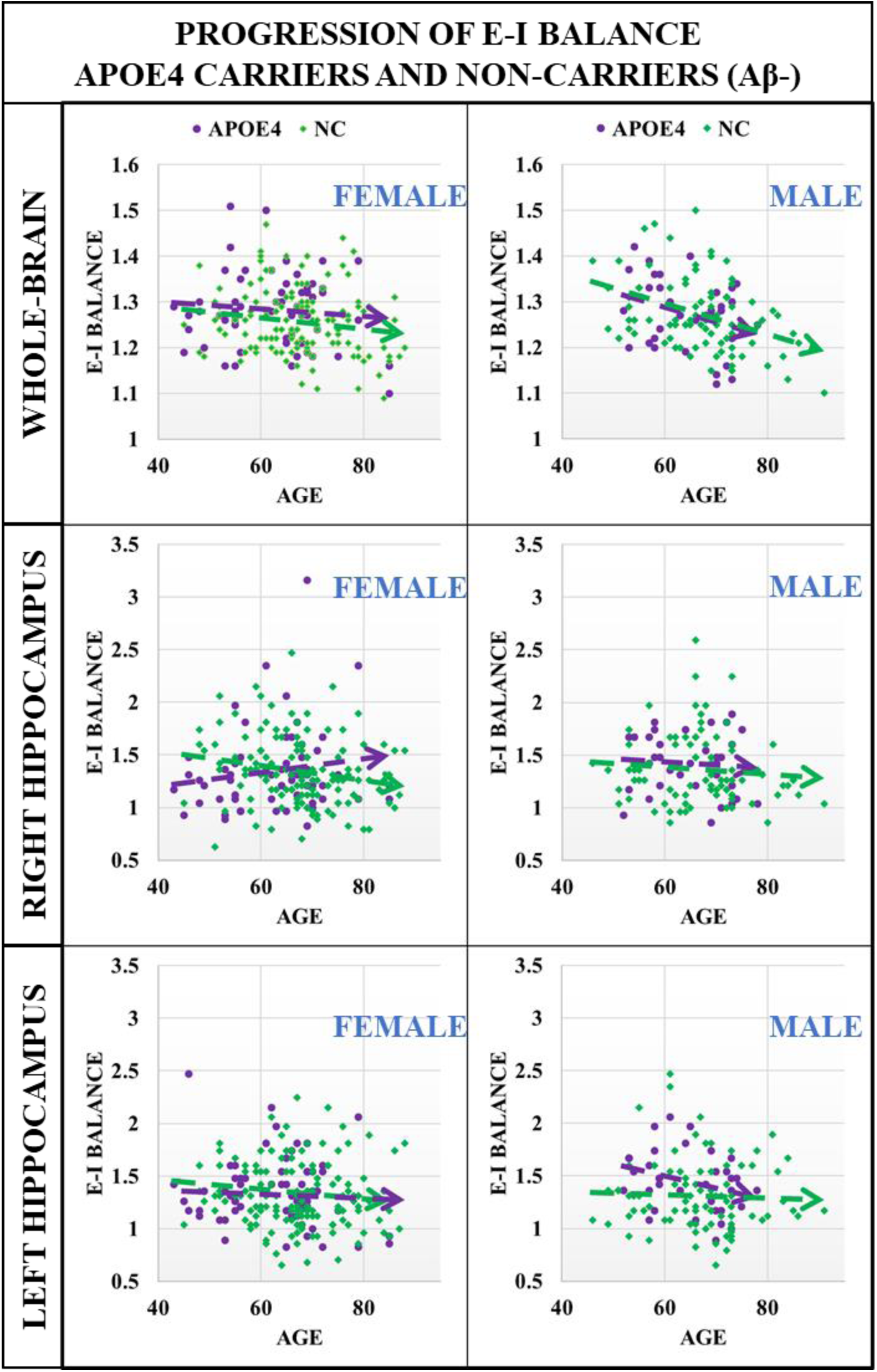
Progression of EIB in cognitively normal males and females, comparing APOE4 and non-carrier (NC) groups that are Aβ-. **Top panel:** whole-brain comparisons for male and female APOE4 carriers and non-carriers (NC) based on linear regression analysis. While there is a group difference in whole-brain EIB between female APOE4 carriers and non-carriers, the difference can be partially explained by age differences in this cross-section, resulting in a p-value of 0.06 for the main effect of APOE4 status. **Middle and Bottom Panels:** Here, only the right hippocampus for females demonstrates divergence for APOE4 carriers as compared to non-carriers. Linear regression modeling to evaluate the interaction of age x APOE4 results in **p = 0.013** (β = 0.014, SE = 0.005). Results suggest that females APOE4 carriers exhibit an increasing effect in EIB in the right hippocampus as compared to non-carriers in the absence of significant amyloid deposits. No significant group differences were identified in the left hippocampus for females or for either side of the hippocampus for men.

### Excitation-inhibition balance in the presence of APOE4 and Amyloid Beta

With the inclusion of individuals with significant Aβ deposits for both APOE4 carriers and non-carriers as described in Table 1, we aim to determine results for three cases in the linear modeling of sex difference in whole brain and hippocampal EIB. First, we evaluate age x APOE interaction, second, we evaluate age x Amyloid interactions, and third we evaluate APOE x Amyloid interactions (noting here that both APOE and amyloid are dichotomous variables). In Figure 3 we present a comparison of hippocampal EIB progression for cognitively normal males and females that are Aβ+. The right hippocampus appears to be significant for both males and females with diverging progression between APOE carriers and non-carriers, however linear regression analysis evaluating interaction effects of age x APOE results in p = 0.012 (β = 0.014, S.E. = 0.012) for the right hippocampus of females only. Regression results are insignificant for males (p > 0.1). Further, linear regression modeling identified a marginally significant interaction effect of age and amyloid status with p = 0.07 (β = -0.02, S.E. = 0.012) for the left hippocampus in females only. No other interaction effects have any significance. Regression results including estimates, p-values, and standard error for all main and interaction effects are shared in Supplemental Table 3. This suggests that the association of EIB with age for both men and women may be affected by amyloid plaques, with more significant differences in female APOE4 carriers.

**Figure 3.**
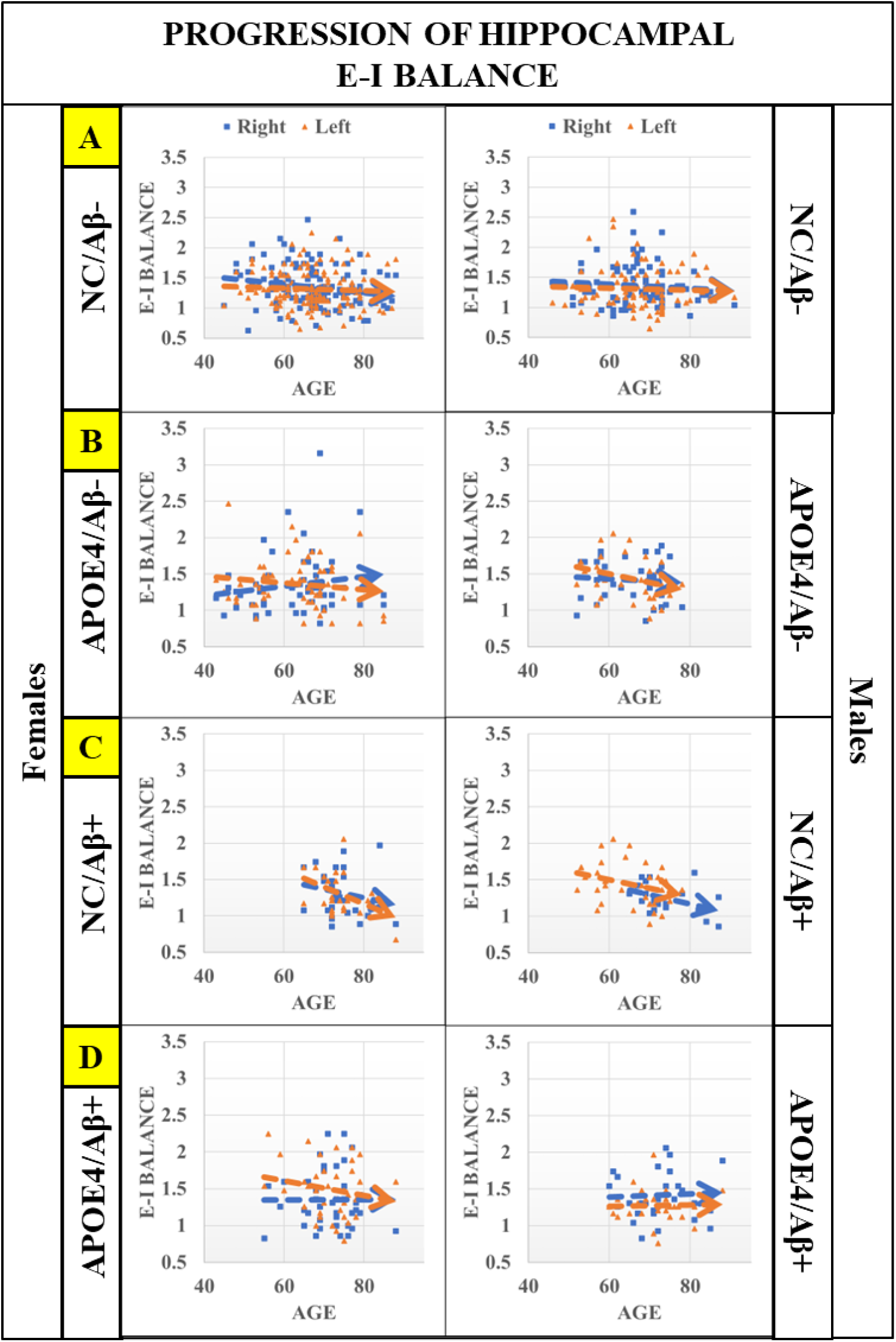
Hippocampal EIB progression for cognitively normal males and females that are Aβ-/Aβ+ (with and without APOE4). **Panels from top to bottom (A-D):** first is a comparison of hippocampal EIB progression in the absence of known AD risk factors (APOE4 and Aβ+), then hippocampal EIB progression for amyloid negative (Aβ-) individuals with at least one APOE4 allele, followed by hippocampal EIB progression of amyloid positive (Aβ+) that are non-carriers (NC) of APOE4, and last is hippocampal EIB progression for individuals that are APOE4 carriers and also Aβ+. These plots suggest that in the presence of significant amyloid (Aβ+), divergence between APOE carriers and non-carriers is significant in the right hippocampus for males and females, however regression modeling to evaluate the interaction effects of age x APOE4 results in **p = 0.012** (β = 0.014, S.E. = 0.012) for females only. Further, linear regression modeling identified a marginally significant interaction effect of age and amyloid status with **p = 0.071** (β = -0.02, S.E. = 0.012) in the left hippocampus for females as well. Regression results are insignificant in the hippocampus (right and left) for males (p > 0.1). Results suggest that features of hyperexcitation (increasing EIB) in hippocampal regions may be driven by APOE4 rather than amyloid for cognitively normal individuals, with more significant impact on females.

### Relationship between excitation-inhibition balance and neuropsychological tests

To evaluate the relationship between EIB for whole-brain dynamics as well as for the hippocampal regions, we perform partial correlation analysis between the neuropsychological test completed in this cohort and EIB of the whole-brain, as well as the left and right hippocampus (for each sex separately). Here we are controlling for primary co-factors of age, APOE4, amyloid status (with the latter two being dichotomous variables), and education due to significant group differences between males and females (shown in table 1). The screening measures of subjects are presented in Table 2, noting that they are a subset of the overall 437 used in this study. Further, as presented in table 2, we note significant differences between males and females for learning memory task I (LM-I), learning memory task II (LM-II), category fluency for vegetables, and the Boston naming test (p < 0.05).

**Table 2.**
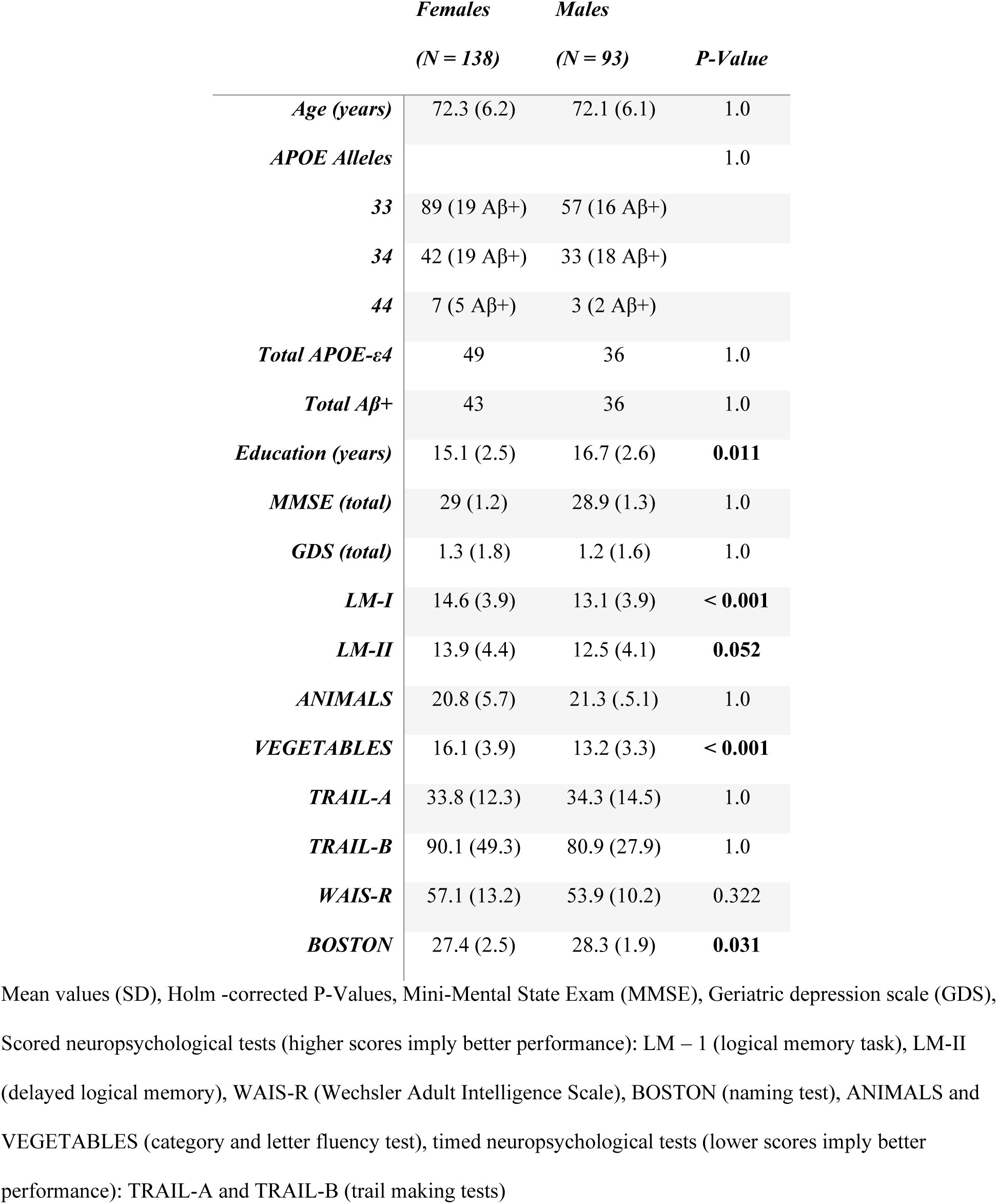
Demographics, and screening measures of participants from OASIS-3 [39] used in secondary analyses involving neuropsychological testing (a subset of those presented in Table 1). All individuals are cognitively normal (CDR = 0) with completed neuropsychological testing

Partial correlation analysis of EIB for the whole-brain, as well as left and right hippocampus with neuropsychological measures are presented in figure 4. Statistically significant correlations are indicated with a star (Holm-corrected P < 0.05), with all results presented in Table 3. The plot in figure 4 highlights differences in both whole-brain and hippocampal relationships with neuropsychological tests in females when controlling for well-known covariates in this dataset related to susceptibility of AD neuropathology. Here, the partial correlation of EIB of the right hippocampus with categorical fluency (ANIMALS) is significant (p < 0.05) and marginally significant with logical memory tests (p < 0.1). Further, the partial correlation of EIB of the whole-brain with trail making test B is significant (p < 0.05) and marginally significant with trail making test A (p < 0.1). We note that the performance in trail making tests is measured in seconds to completion, hence shorter times imply better performance. Both sets of results are specific to the female group. Partial correlations of EIB with neuropsychological tests are insignificant for the male group when controlling for co-factors of age, APOE, and amyloid status. Further, there is a more right hemispherical dominance in terms of hippocampal EIB for females compared to left hemispherical dominance for males in most tasks, however males do not have any statistically significant correlations between EIB of the hippocampus or whole-brain with these neuropsychological tests when controlling for co-factors of age, APOE, and amyloid status, and education. This, in contrast to females that do have more significant relationships with the right hippocampus. We note here the pattern of Trail making tests having more influence from whole-brain dynamics than either hippocampal region for both males and females (albeit more significantly for the latter).

**Figure 4.**
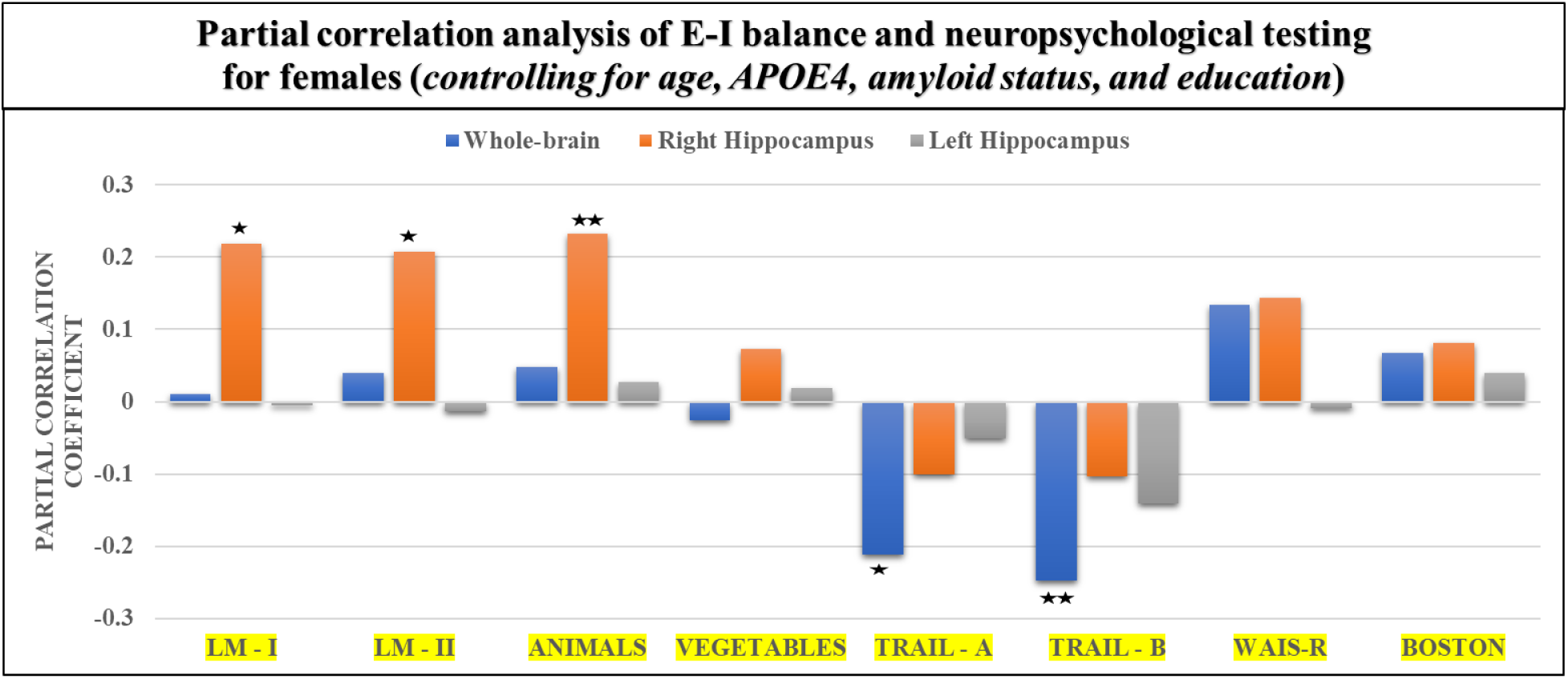
Partial correlation analysis of EIB (whole-brain, left and right hippocampus) and neuropsychological testing for females, controlling for co-variates of age, APOE4, and amyloid status. Presented here are partial correlation analysis of EIB for the whole-brain, as well as left and right hippocampus with neuropsychological measures. Statistically significant correlations are indicated with two stars (Holm-corrected p < 0.05) and partially significant correlations are indicated with a single star (Holm-corrected p < 0.1). There are no statistically significant (or marginally significant) results for males, and therefore are not included in this figure (all results for both sexes included in Table 3). The plot here highlights differences in both whole-brain and hippocampal relationships with neuropsychological tests when controlling for well-known covariates in this cognitively normal cross-section in terms of susceptibility to AD neuropathology. Here, performance in category fluency tests (ANIMALS) demonstrates a significant positive correlation of the right hippocampus, while performance in trail making test B is significantly correlated with the EIB of the whole-brain. We further note that logical memory tests (I, II) have marginally significant positive partial correlations with right hippocampus EIB as well, while trail making test A is also marginally significant with whole-brain EIB. We note that for trail-making tests, performance is determined by the shortest time to completion, so negative correlations indicate a positive association with performance. Further, visually we see a hemispherical dominance in terms of right hippocampal EIB with neuropsychological test performance. We note here the pattern of Trail making tests having more influence from whole-brain dynamics than either hippocampal region for both males and females (albeit more significantly for the latter). In sum, results suggest that increasing EIB of the hippocampus seen in females, particularly those at increased risk of developing AD, correlated with improved performance in neuropsychological tests for females.

**Table 3.** Partial correlation analyses of EIB and neuropsychological testing, controlling for the effects of age, sex, APOE4, and amyloid status.

|  | *Whole-Brain* |  | *Hippocampus (right)* |  | *Hippocampus (left)* |  |
| --- | --- | --- | --- | --- | --- | --- |
|  | Females | Males | Females | Males | Females | Males |
| ***LM – I*** | 0.011 (0.9) | -0.084 (1.0) | **0.22 (0.078\*)** | -0.043 (1.0) | -0.004 (0.965) | -0.261 (0.1) |
| ***LM – II*** | 0.041 (1.0) | -0.047 (0.66) | **0.21 (0.097\*)** | -0.063 (1.0) | -0.013 (1.0) | -0.21 (0.213) |
| ***ANIMALS*** | 0.048 (1.0) | -0.082 (1.0) | **0.23 (0.048\*\*)** | -0.029 (1.0) | 0.027 (1.0) | -0.161 (0.517) |
| ***VEGETABLES*** | -0.026 (1.0) | 0.133 (1.0) | 0.072 (0.413) | 0.112 (1.0) | 0.019 (1.0) | 0.243 (0.152) |
| ***TRAIL - A*** | **-0.21 (0.098\*)** | -0.184 (0.652) | -0.1 (0.75) | -0.112 (1.0) | -0.051 (1.0) | 0.062 (1.0) |
| ***TRAIL - B*** | **-0.252 (0.034\*\*)** | -0.113 (1.0) | -0.1 (0.951) | -0.112 (1.0) | -0.144 (0.855) | -0.016 (0.886) |
| ***WAIS-R*** | 0.133 (0.731) | 0.061 (1.0) | 0.142 (0.49) | 0.025 (0.821) | -0.009 (1.0) | -0.1 (1.0) |
| ***BOSTON*** | 0.067 (1.0) | 0.182 (0.721) | 0.082 (0.662) | 0.133 (1.0) | 0.041 (1.0) | 0.252 (0.148) |

Linear partial correlation coefficient $\beta$ (Holm-corrected P-Values), scored neuropsychological tests (higher scores
imply better performance): LM – I (logical memory task), LM-II (delayed logical memory test), WAIS-R (Wechsler
Adult Intelligence Scale), BOSTON (naming test), ANIMALS and VEGETABLES (category and letter fluency
test), timed neuropsychological tests (lower scores imply better performance): TRAIL-A and TRAIL-B (trail making tests),
\*: Adjusted $P < 0.1$
\*\*: Adjusted $P < 0.05$

## Discussion

The present study was conducted to determine how excitation-inhibition balance (EIB) in cognitively normal individuals may be associated or influenced by AD risk factors such as age, sex, APOE genotype, and Aβ. Using a framework based on statistical physics, we computationally inferred EIB to macroscopically evaluate whole brain and hippocampal signatures of hyperexcitation. The novelty of this model lies in integrating microscale principles at a connectome level to bridge the gap between cell-to-network level degeneration. We previously demonstrated age- and sex-dependent hyper-excitation within regions of the default mode network (DMN) in a cohort of Aβ-cognitively normal female carriers of APOE4 allele [12,59] where traditional measures of functional connectivity were unable to detect the subtle differences in brain dynamics between female carriers and non-carriers in middle age [60,61]. Several studies have investigated EIB as an effect of the normal aging and neurodevelopmental process [3,62–64], however few incorporate known sex-differences in brain activity and development [65].

The results presented herein demonstrated that in the absence of AD risk factors (APOE-ɛ4/Aβ+), whole-brain EIB progresses differently between males and females with age, with males decreasing more significantly as they get older. Further, performing regression modeling with the inclusion of individuals having at least one APOE-ɛ4 allele (Aβ-) produced a significant age-by-APOE interaction in the right hippocampus for females only. The effect remained unchanged with inclusion of Aβ+ individuals. Last, we performed partial correlation analyses between EIB and neuropsychological testing, revealing positive associations between hippocampal and whole-brain EIB with test performance (for females only). Significant relationships between EIB in the right hippocampus with categorical fluency and whole-brain EIB with a trail-making test were identified. Each of these results will be discussed below.

The investigation in the context of normal aging is based on N = 228 (143 Female) individuals that have the APOE ε3/ε3 combination and classified as Aβ-. We note as mentioned in the methods section, the exclusion of individuals with the APOE-ε2 allele. Deciding whether to include or exclude subjects with additional allele combinations including APOE-E2 merits further discussion. In this study, we aimed to expand on our previous work in a larger cohort evaluating EIB as a potential MRI-based biomarker demonstrating pathophysiological differences in males and females that may confer latter greater vulnerability to AD neuropathology. By including individuals with additional allele combinations, we would then have to include an additional variable in our analysis, namely allele pair combinations. While it may provide additional insight into an EIB spectrum among different allele combinations, it would also decrease the statistical power of our results as some allele combinations would have significantly fewer subjects than others for each sex (even more so when breaking down by amyloid positivity as well). Our future work will aim to drill down further, beyond simple ε3-ε4 comparisons to better understand the relationship between the allele combinations and EIB. Here, our study demonstrates that alterations in the excitation/inhibition balance (EIB) are complex mechanisms that are associated with aging, sex, and the APOE4 genotype before cognitive impairment occurs. Interestingly, we observed a reduction in EIB with age in males, presented in Figure 1, which may be protective against hyperexcitability. It has been reported that alterations of tonic inhibition lead to hyperexcitability, which could contribute to the development of neurodegenerative diseases [66]. Thus, the reduction of excitation with aging in males might serve as a protective mechanism against neurodegeneration. Further research is needed to determine the underlying mechanisms of this phenomenon and its implications for disease progression. We acknowledge that it is difficult to interpret whether females have increased excitatory tone or males have increased inhibitory tone with respect to healthy aging and what other factors may be involved (genetic, hormonal, physiological, etc.).

These sex differences are highlighted further with the inclusion of individuals with APOE-ε4 and provide additional support to our previous findings in that females with at least one APOE4 allele are at higher risk of AD neuropathology due to a hyperexcitation of critical brain regions. Specifically, we found that EIB diverges significantly in the right hippocampus for female carriers, with no change in either side of the hippocampus for males. Shown in Figure 2, these findings are biologically significant in the following two important ways:

1. They are consistent with the recently proposed theory of a hyper-excitable state in the earliest AD pathology, due to the susceptibility of inhibitory GABAergic interneurons to ApoΕ4 fragment-mediated neurotoxicity (in the dentate gyrus or DG of hippocampus).
2. They may help explain sex differences in AD (females are at a substantially greater risk of developing AD) by linking the earliest pre-clinical changes of AD pathology, i.e., hyper-excitability in the hippocampus, to reported sex-differences in functional connectivity developed in early adulthood and sustained across the life span. Indeed, fMRI studies have shown that females exhibit higher resting-state functional connectivity in the temporal lobe, and thus likely a differentially higher excitation relative to inhibition). Given the complexity multifactorial nature of AD, network-level neuroimaging metrics may be more sensitive to subtle structural and functional disruptions than traditional analyses [67,68].

In fact, our results suggest that APOE, but not amyloid, may be the driver for increased EIB deviations as demonstrated by the interaction effect of APOE and age, having no change in statistical significance with the inclusion of Aβ+ individuals. However, in this cohort (OASIS-3), the cognitively normal group is heavily skewed in terms of Aβ-individuals, which may result in low statistical power for differentiating between APOE and amyloid-mediated effects in this cross-section. In our analysis, we recognize several limitations that may impact the interpretation of our findings. An age-matched cohort was not included in our comparisons, an aspect which may have influenced our results and interpretation., Inclusion of an age-matched cohort in the future may allow for a more robust comparison and analysis of an APOE4 effect. Further, our study lacks information regarding tau load, preventing the study of EIB with relation to the triad of AD risk (APOE4, amyloid, and tau) in terms of healthy aging and AD risk. Some studies have found that the interaction effect between APOE4 and amyloid-β, rather than the sum of their independent effects, was related to increased tau load in Alzheimer’s disease-vulnerable regions [69,70]. This may help to explain the lack of interaction between APOE4 and amyloid-β. The interaction between one APOE4 allele and amyloid-β is hypothesized to be related to increased tau load, while the interaction between amyloid-β and two APOE4 alleles was related to a more widespread pattern of tau aggregation [69,71]. With Aβ-mediated hyper-excitation emerging as a promising candidate mechanism of AD pathogenesis: inhibitory GABAergic dysfunction is predictive of increased neuronal activity, while the increased damage of dendrites may be associated with a silencing effect of neuronal circuits (the yin and the yang of AD degeneration). However, recent evidence however, points to hyper-excitability of neuronal circuits as a precursor to the formation of amyloid plaques [70,72,73], and a major driver of the cascading effect of tau pathology, propagating from the entorhinal cortex to the hippocampus and beyond to other cortical regions [74]. In fact, one study has even demonstrated differential effects of sex by APOE interaction on tau accumulation, in that only one APOE ε4 allele is sufficient to increase tau accumulation in females, while two APOE ε4 alleles are needed to cause a similar effect in males [75]. In future studies, we aim to identify EIB changes which may be related to APOE, Aβ, and tau in healthy aging as well as in individuals that are cognitively impaired and/or have AD to identify a more complete picture of EIB progression in relation to neurodegeneration and these risk factors. The complex, multifactorial nature of Alzheimer’s disease makes it essential that we continue to develop techniques to meaningfully integrate multiple neuroimaging modalities to increase the sensitivity to detect very early network aberrations associated with risk for this disease.

Last, we found an association between EIB and neuropsychological testing, specifically in females when controlling for age, education, APOE, and amyloid positivity. We note that while it is true that amyloid presence is a defining feature of AD pathology, it is also a risk factor that may precede the clinical manifestations of the disease. Controlling for amyloid status allows us to account for the influence of preclinical AD pathology on the observed associations between EIB and neuropsychological variables. Further, including both APOE and amyloid in the study of hyperexcitation differences in cognitively normal individuals allows us to account for potential genetic and pathological influences on neural activity, as both factors have been implicated in the development and progression of Alzheimer’s disease.

As shown in figure 4, partial correlation analyses revealed positive associations between EIB and neuropsychological testing for females, having significant relationships between EIB in the right hippocampus with categorical fluency and whole-brain EIB with a trail-making test. Performance in category fluency tests (ANIMALS) is significantly correlated with EIB of the right hippocampus for women, while performance in trail making test B is significantly correlated with the EIB of the whole-brain. We further note that logical memory tests (I, II) have marginally significant partial correlations with EIB of the right hippocampus as well, while trail making test A is also marginally significant with whole-brain EIB. We should clarify that the partial correlations we’ve presented highlight the unique contribution of EIB to neuropsychological performance, independent of the influence of covariates such as age, education, APOE, and amyloid status. For example, a lower partial correlation between left hippocampal EIB and verbal memory tasks does not necessarily signify a negligible overall association. The traditionally acknowledged association between left Hippocampus and verbal memory could indeed be a downstream consequence of factors such as APOE and amyloid.

Further, there is also a more hemispherical dominance in terms of hippocampal EIB however males have no significant correlations in the EIB as compared to females that have more significant relationships with the right hippocampus. The observed association between right hippocampal EIB and verbal memory tasks, as opposed to left hippocampal (left) EIB, might appear surprising given the traditional view of left Hippocampus being more involved in verbal memory processing. However, recent research has suggested that the lateralization of memory functions is more complex than initially thought, with some studies showing the involvement of both sides of the hippocampus in verbal memory tasks [76,77]. To that end, these results shed light on a pattern of hemispheric dominance in the hippocampus, with women demonstrating a stronger correlation with the right hippocampal EIB. This finding might suggest sex-specific patterns of hippocampal involvement within the context of AD risk, a potentially important consideration when assessing the cognitive status and potential AD risk of female patients. Further research is needed to confirm this hypothesis and better understand the underlying mechanisms. We note here the pattern of Trail making tests having more influence from whole-brain dynamics than either hippocampal region for both males and females (albeit more significantly for the latter). The pattern of Trail Making Tests having more influence from whole-brain dynamics than hippocampal regions for both males and females suggests that these tests might be more sensitive to global cognitive changes associated with AD risk, rather than being specific to hippocampal function [78]. This further highlights the importance of considering sex-specific differences when evaluating the relationship between EIB and cognitive performance in the context of AD risk. Further, while our analysis does control for factors like APOE and amyloid status, it’s crucial to note that our intention is not to dismiss the role of AD risk factors. Instead, we aim to delve deeper into the independent role of EIB in cognitive performance, beyond these known risk factors.

These results suggest that increasing EIB is correlated with improved performance in neuropsychological tests for females and related two ongoing hypothesis; the first being that increased excitation (a positive shift in EIB) is associated with better performance in neuropsychiatric testing (as others have demonstrated with transcranial magnetic stimulation and transcranial direct current stimulation [79,80]), and second being that potential hyperexcitation due to APOE-E4 both precedes and may even influence increased spread of amyloid plaques before cognitive impairment is identified [81]. It is also possible that the APOE-E4 genotype may be protective against a subtle deficit associated with β-amyloid pathology, and consistent with the antagonistic pleiotropy hypothesis; whereby a gene controls both beneficial and detrimental traits – and provides novel evidence that these effects persist into older age, even among individuals who cognitively unimpaired (but may have additional AD risk factors) [82–84]. Despite strong evidence of the clinical utility of neuropsychological tests, however, it is unclear what neurobiological information the test scores provide. While the progression of the pathophysiological process of dementia strongly affects neuropsychological function [85,86], a large amount of variance is left unexplained even with multimodal biomarker information [87–89]. Moreover, it is hard to predict whether the presence of pathophysiological markers directly leads to neural dysfunction within the behaviorally relevant regions [90].

Further, the cross-sectional design of our study precludes us from making causal inferences about the relationship between EIB and Alzheimer’s disease (AD) neuropathology. Longitudinal analysis would be necessary to investigate EIB as a predictor of individuals who subsequently develop AD neuropathology. In addition, neuropsychological test data was only available for individuals aged 65 or older, which may limit the generalizability of our findings to younger populations. Last, we recognize that hormonal differences between males and females could be a confounding factor in our study, as they may influence EIB and susceptibility to AD. Future research should consider hormonal information when investigating sex differences in EIB and its relation to neurodegenerative diseases (in particular, evaluating female EIB progression separated by pre - and post-menopause).

Last, our findings contribute to the understanding of how EIB is involved in neurodevelopmental disorders and neurodegenerative diseases (such as autism spectrum disorder, schizophrenia, attention deficit/hyperactivity disorder and Alzheimer’s disease) and how sex differences in EIB may be leveraged to inform therapeutic intervention and clinical endpoints. As a therapeutic target, transcranial direct current stimulation (tDCS) has emerged as a promising therapy for rehabilitating neurodevelopmental disorders such as autism, schizophrenia, attention deficit/hyperactivity disorder, and even epilepsy [80,91–93]. The mechanism behind tDCS’s effects is the modulation of excitatory and inhibitory activity, making it a valuable tool for restoring excitation-inhibition balance. Clinical studies have demonstrated that tDCS therapy is well-tolerated by patients and appears to improve behavior and cognitive functions. Further, AD patients present an increased prevalence of epilepsy, with neural hypersynchrony and aberrant oscillatory activity contributing to cognitive deterioration [94–96]. Levetiracetam (LEV) has shown effectiveness in reducing epileptiform activity in AD patients, correlating with improved cognitive performance [97,98]. Ongoing clinical trials are assessing LEV’s effectiveness in AD patients without epilepsy, indicating potential future clinical applications of our hybrid connectome and EIB in the context of epilepsy and Alzheimer’s disease [99].

Our results, along with the growing body of evidence conceptualizing AD neuropathology in terms of an imbalance of excitation and inhibition suggests that females early in the disease progression (particularly those with the APOE-E4 genotype) may be more responsive to tDCS or low doses of anticonvulsants. The hybrid connectome (rs-SC) framework used to infer excitation-inhibition balance at an individual level, may be a step towards an MRI-based biomarker to identify hyperexcitation-induced neurodegeneration before traditional indicators of AD arise and potentially evaluate therapeutic interventions and clinical endpoints.

## Acknowledgements

The magnetic resonance imaging (MRI) and neuropsychological tests data that support the findings of this study are available in [“OASIS-3”] with the identifier(s) [https://doi.org/10.1101/2019.12.13.19014902] [39]. OASIS-3: Longitudinal Multimodal Neuroimaging: Principal Investigators: T. Benzinger, D. Marcus, J. Morris; NIH P30 AG066444, P50 AG00561, P30 NS09857781, P01 AG026276, P01 AG003991, R01 AG043434, UL1 TR000448, R01 EB009352. AV-45 doses were provided by Avid Radiopharmaceuticals, a wholly owned subsidiary of Eli Lilly. OASIS-3 data are openly available to the scientific community at https://www.oasisbrains.org. Prior to accessing the data, users are required to agree to the OASIS data use terms (DUT), which follow the creative commons attribution 4.0 license.

## Data Availability

The data supporting the findings of this study are available on request from the corresponding author. The data are not publicly available due to privacy or ethical restrictions.

## Supplementary Material

### MRI data preprocessing

First, extra-cerebral tissue was removed from the T1-weighted anatomical scans using a robust automated brain extraction program trained on manually “skull-stripped” MRI data (ROBEX) [100]. Skull-stripped volumes were visually inspected, and manually edited if needed. Intensity inhomogeneity normalization using the MNI nu_correct tool (www.bic.mni.mcgill.ca/software/) was performed on the anatomical scans. For diffusion-weighted MRI (DWI) data, we use FSL eddy-correct tool (http://www.fmrib.ox.ac.uk/fsl) to correct for head motion and eddy current distortions, each subject’s raw DWI volumes were aligned to the b_0_ image, and the gradient table modified accordingly. Using the Brain Extraction Tool (BET) from FSL [101], non-brain tissue was removed from the DWIs. And lastly, we adopt dtifit function from FSL to compute fractional anisotropy (FA) map for each subject.

### Whole brain tractography

For computing whole-brain tractography, we leveraged the Runge-Kutta (RK2) algorithm, which uses the second order Runge–Kutta method to solve a differential equation to estimate the fiber trajectory more reliably [102]. The main mathematical technique behind RK2 is to equate the tangent vector with the principal eigenvector. The Diffusion toolkit (http://trackvis.org/dtk/) was used to implement the RK2 approach. Further, fiber tracking was restricted to regions where FA ≥ 0.18, thus avoiding grey matter (GM) and cerebrospinal fluid (CSF). Note that fiber paths were stopped if the fiber direction encountered a sharp turn (with a critical angle threshold ≥ 45°). These parameter settings for each method that had been previously optimized by our group [103] or others; in most cases, they were the default parameter settings of the methods. To avoid undue complexity, we concede that changing these parameters could conceivably affect how the methods are ranked. Moreover, all fibers shorter than 10 mm were filtered out, as these were much more likely to be false positive fibers. During the processing stage of the imaging sessions a further 18 sessions were excluded from this study due to poor structural imaging.

### Computing structural networks

Brain regions (ROIs) covering the entire brain comprised 132 regions (91 cortical ROIs from the FSL Harvard-Oxford Atlas maximum likelihood cortical atlas, 15 subcortical ROIs from the FSL Harvard-Oxford Atlas maximum likelihood subcortical atlas [104] and 26 cerebellar ROIs from the AAL atlas [105] which are included in the Conn toolbox. Midline cortical masks were bisected to separate left and right components so as to define hemispheric ROIs for each cortical region where possible. As this is a probabilistic atlas, masks were thresholded at 10% to ensure inclusion of tissue along the gray-WM interface, where fiber orientation mapping and tractography are most reliable [103]. In order to map these 132 ROIs from atlas space to subject’s DWI space, we firstly map the atlas T1 to the subject T1 using a correlation ratio cost function and map a subject’s T1 to subject’s FA using a mutual information cost function. These two steps can be done using linear registration (i.e., flirt[101] function in FSL toolbox) as described in the processing steps. Last, after combining transformation matrices from previous steps, the 132 ROIs were transformed to each subject’s DWI or FA space using nearest-neighbor interpolation. Each voxel was uniquely assigned to the mask for which it had the highest probability of membership, thus ensuring that ROI masks did not overlap with each other after registration.

The number of detected fibers connecting brain regions was determined from the tractography results. If a fiber intersected two ROIs, it was considered as connection between them. This process was repeated for all ROI pairs, to compute a whole-brain fiber connectivity matrix. This matrix is symmetric, by definition, and has a zero diagonal (no self-connections [106]). In this study, to avoid computation bias when constructing the hybrid resting-state structural connectome, each brain matrix was normalized by the maximum value in the matrix, as matrices derived from different subjects may have different scales and ranges.

### Functional MRI processing and functional network construction

The raw fMRI signals were processed using the CONN toolbox (McGovern Institute for Brain Research, MIT, USA) running under MATLAB (Mathworks), and FSL (FMRIB Software Library v5.0, http://www.fmrib.ox.ac.uk/fslwiki, Analysis Group, FMRIB, Oxford, UK). For a detailed description of the CONN pipeline, the online documentation can be consulted (https://web.conn-toolbox.org/home). Here a short summary is provided for the main steps of functional realignment and unwarping, slice-timing correction, outlier identification, direct segmentation and normalization, functional smoothing, followed by regressing out confound effects.

First, a realigning and unwarping procedure is used for co-registration and resampling of all scans to the first scan using b-spline interpolation [107]. This is followed by temporal misalignment correction by time-shifting and resampling the slices using the sinc-interpolation (this matches the time in the middle of each acquisition) [108]. Next, the outlier identification procedure is followed, with the individual volumes flagged as potential outliers if the framewise displacement (FD) of a given volume is above the default CONN Toolbox threshold of 0.9 mm or global BOLD signal changes above five standard deviations. Once potential outliers are removed, averaging the remaining volumes determines the new reference image. This image is then used for segmentation and normalization. All samples must then be normalized into standard MNI space. Scans are segmented into white matter (WM), gray matter (GM), and cerebrospinal fluid (CSF). In this step, the structural T1 images are used to improve the quality of the registration, using the unified segmentation and normalization procedure [109]. To increase the signal-to-noise ratio, fMRI signals are smoothed by convolving them with an 8 mm Gaussian kernel. Last, confound effects from motion artifact, WM, and CSF were regressed out of the signal [110]. This pipeline has been applied and verified in our previous studies [59,60,110]. In this study, the completeness of the fMRI imaging is an important component for our methodology. Thus, after evaluating the processed fMRI imaging results, a further 26 imaging sessions were removed from this study for missing more than fifty percent of fMRI volumes (i.e., less than 82 points in the time series). Once the resting-state fMRI signal is preprocessed, functional brain networks can be derived by computing Pearson correlations between all pairs of ROIs in the BOLD signal time series using the same 132 labels as in the structural brain networks. Using the same 132 labels as in the structural brain networks, functional brain networks were derived by computing Pearson correlations between all pairs of ROIs in the BOLD signal time series. A subset of 105 regions was used in this study (after computing the functional networks, a subnetwork was extracted excluding the brain stem and cerebellar regions).

### Constructing a metric of excitation-inhibition balance (EIB) based hybrid resting-state structural connectivity

Recently our team [12,56] combined MRI-derived multimodal connectome data (resting-state functional and structural) with statistical physics to obtain a macroscopic account of inhibitory and excitatory long-range connections. Here, we employed the Ising model, a well-known spin-glass model from statistical physics in which the states, also referred to as “spin configurations”, of interacting units – in our case brain regions connected by white matter edges – are constrained to be either 1 (“active”) or -1 (“inactive”). As described in [12,56] we construct a function-by-structure embedding (FSE) using a constrained pseudolikelihood estimation technique wherein pairwise interaction coefficients are inferred from the observed data (BOLD time series). The model assumes binary data, hence we binarize the resting-state fMRI signals. The binarized activity pattern of all ROIs at time t (t = 1,2, …, t_max_;) is denoted **S**(t) = s_1_(t), s_2_(t), … s_N_(t) ɛ {−1, +1}^N^. Note that tmax is determined as a result of the fMRI scan time. Here s_1_(t) = ± 1 indicates that an ROI is either active (+1) or inactive (-1). First, the time series goes through a z-score normalization procedure, resulting in zero mean and unitary variance. The interaction J_i,j_ between two regions should be directly linked back to the diffusion MRI-derived structural connectivity between them as informed by tractography, so we add a constraint to the Hamiltonian function as H(s) = − ∑_i<j_ J_i,j_s_i_s_j_, such that |J_i,j_| ∝ W_i,j_, where W_i,j_ is the structural connectivity between pairs of ROIs, and the external force or bias terms are dropped in the case of resting-state. This ensures that in the pseudolikelihood estimation of **J**, we constrain it with the structural connectivity (under the assumption that structural connectivity informs spin models governing brain dynamics). Conceptually, if two regions are co-active or co-inactive, then the interaction (J_i,j_) is likely positive (excitatory), and if one region is active while the other is inactive, then the interaction (J_i,j_) is likely negative (inhibitory).

Hence, the framework for constructing the hybrid resting-state structural connectome (rs-SC) using the FSE framework follows a 3-stage processes (described in more detail in our previous work) [12,56]. In stage 1, we estimate both edge strength in the connectome network as well as a sign (+/-) representing excitatory or inhibitory interactions (J_i,j_) based on structural and functional MRI imaging. In stage 2, two metrics are used to evaluate parameter quality, namely a similarity metric **S**_**m**_ and the maximum of a functional correlation function max **f**_**c**_. The similarity metric S_m_(**J**) is simply the correlation between |rs-SC| and the structural connectome. This metric is used to gauge the quality of the constraint component in the framework. Further we generate a correlation function f_c_(FSE) by simulating the Ising model with MCMC simulations to reconstruct FC from the inferred interactions between brain regions (J_i,j_), computing a Pearson correlation between observed and simulated functional connectivity. Here we identify the max f_c_(**J**), which represents the maximum achieved correlation between observed and reconstructed FC under empirically determined FSE parameters [12,56]. As these values are dependent on parameter choices within the FSE framework, in Stage 3 a grid search is performed to find the optimal values for the tuned parameters which maximizes f(FSE) = max **f**_**c**_ + **S**_**m**_ which intends to take equal weight between the underlying structure of the rs-SC and the accuracy with which it can reconstruct FC These two metrics were computed for each subject individually, to identify the optimal parameters for constructing a subject-level hybrid resting-state structural connectome (rs-SC). The code used to compute the rs-SC can be found at https://github.com/iforte2/hybrid-connectome. This procedure is followed for all subjects in constructing an optimized J matrix per subject, which we term the resting-state structural connectome or rs-SC. In this manner, dynamics generated using this J matrix as the interaction coefficients of an Ising spin-glass system maximally explain the observed resting-state fMRI time series.

Note that although our framework for constructing the rs-SC does not explicitly model brain activity at a neuronal level, several seminal studies have used the Ising model to investigate neuronal firing patterns with success, showing that ensemble level activities can be well explained by pairwise Ising interactions [24,30]. To be clear, from a connectomics perspective, if several brain regions are identified to have an increase in positive edges in the rs-SC, collectively, that would suggest a wider-spread pattern of coupling (i.e., more likely to exhibit a pattern of global coupling) that may subserve hyperexcitation. It is in this context that we conceptualize the Excitation-Inhibition (E-I) ratio, a global (whole-brain) or local (ROI-level) estimation of EIB, computed as the sum of positive edges divided by the sum of negative edges. For example, if a brain region in the network has 45 positive interactions (edges) and 34 negative interactions with other brain regions, then the 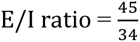, or 1.32 (a value of 1 indicates perfect EIB).

---

**Supplemental Figure 1.**
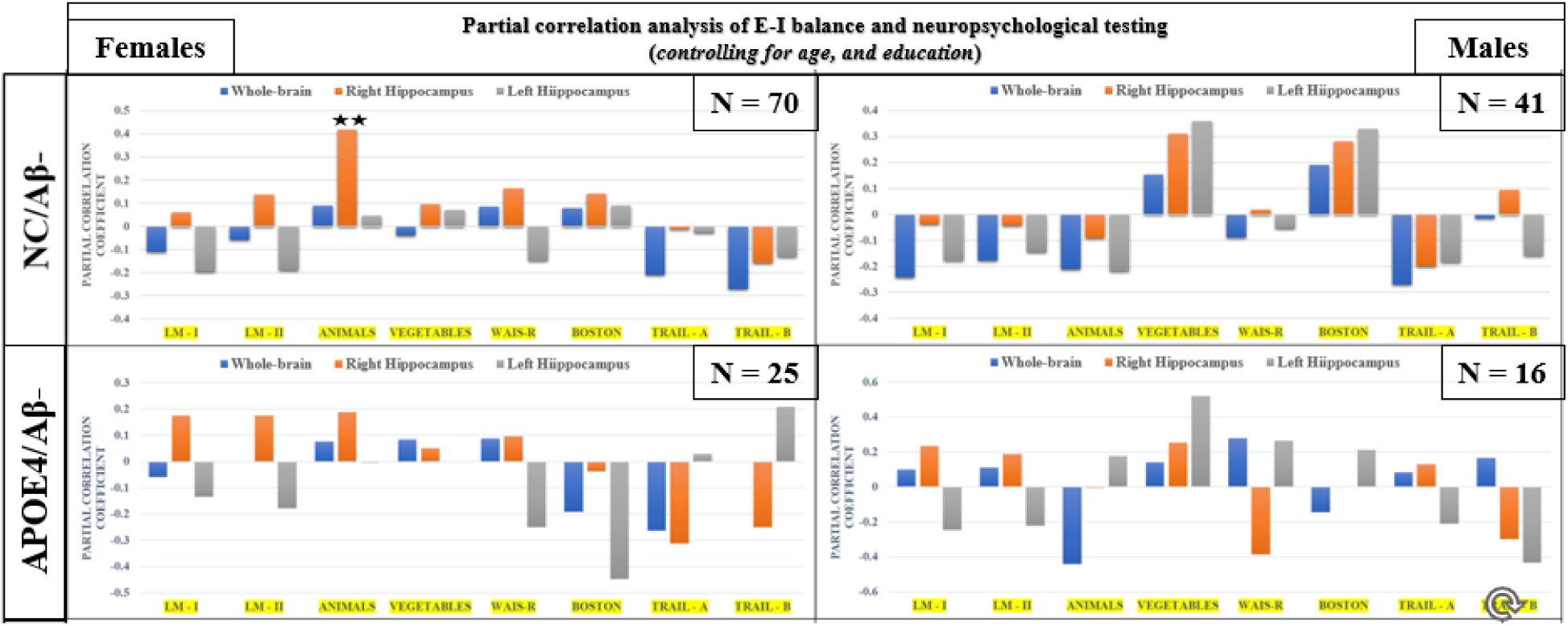
Partial correlation analysis of EIB (whole-brain, left and right hippocampus) and neuropsychological testing for females, controlling for co-variates of age and education. Presented here are partial correlation analysis of EIB for the whole-brain, as well as the hippocampus (right and left) with neuropsychological measures. Statistically significant correlations are indicated with two stars (Holm-corrected p < 0.05).

**Supplemental Table 1.** Regression model results examining EIB as a function of age and sex interactions for cognitively normal males and females without any known predispositions to AD neuropathology (i.e., individuals with CDR = 0, Aβ-, and APOE-ε3/ε3 allele combinations).

|  | *Whole-Brain* |  |  | *Hippocampus (RH)* |  |  | *Hippocampus (LH)* |  |  |
| --- | --- | --- | --- | --- | --- | --- | --- | --- | --- |
| | $\beta$ | P | S.E. | $\beta$ | P | S.E. | $\beta$ | P | S.E. |
| *Intercept* | 1.25 | 0 | 5.90E-03 | 1.347 | 0 | 0.026 | 1.315 | 0 | 0.026 |
| *Age* | -0.001 | 0.057 | 0.0006 | -0.007 | 0.018 | 0.002 | -0.002 | 0.496 | 0.002 |
| *Sex* | 0.015 | 0.102 | 0.009 | 0.014 | 0.735 | 0.043 | -0.004 | 0.919 | 0.043 |
| *Age:Sex* | -0.002 | 0.046 | 0.001 | 0.003 | 0.461 | 0.004 | 0.0004 | 0.936 | 0.004 |

**Supplemental Table 2.** Regression model results assessing EIB as a function of age and APOE4 interactions for individuals that are Aβ- to evaluate the influence of the APOE4 allele on EIB in healthy aging.

|  | *Whole-Brain* |  |  | *Hippocampus (RH)* |  |  | *Hippocampus (LH)* |  |  |
| --- | --- | --- | --- | --- | --- | --- | --- | --- | --- |
| | $\beta$ | P | S.E. | $\beta$ | P | S.E. | $\beta$ | P | S.E. |
| *Intercept* | 1.257 | 0 | 0.006 | 1.354 | 0 | 0.027 | 1.317 | 0 | 0.026 |
| *Age* | -0.001 | 0.065 | 0.0007 | -0.007 | 0.022 | 0.003 | -0.002 | 0.493 | 0.002 |
| *Sex* | 0.018 | 0.071 | 0.01 | 0.011 | 0.809 | 0.045 | -0.004 | 0.912 | 0.043 |
| *APOE4* | 0.022 | 0.063 | 0.011 | 0.015 | 0.769 | 0.053 | 0.029 | 0.566 | 0.051 |
| *Age:Sex* | -0.002 | 0.053 | 0.001 | 0.003 | 0.478 | 0.004 | 0.0004 | 0.935 | 0.004 |
| *Age;APOE4* | 0.0004 | 0.730 | 0.001 | 0.013 | 0.012 | 0.005 | -0.002 | 0.602 | 0.005 |
| *Sex:APOE4* | -0.029 | 0.117 | 0.019 | 0.034 | 0.692 | 0.086 | 0.088 | 0.286 | 0.082 |
| *Age:Sex:APOE4* | -0.000 | 0.881 | 0.002 | -0.013 | 0.199 | 0.010 | -0.007 | 0.443 | 0.009 |

**Supplemental Table 3.**
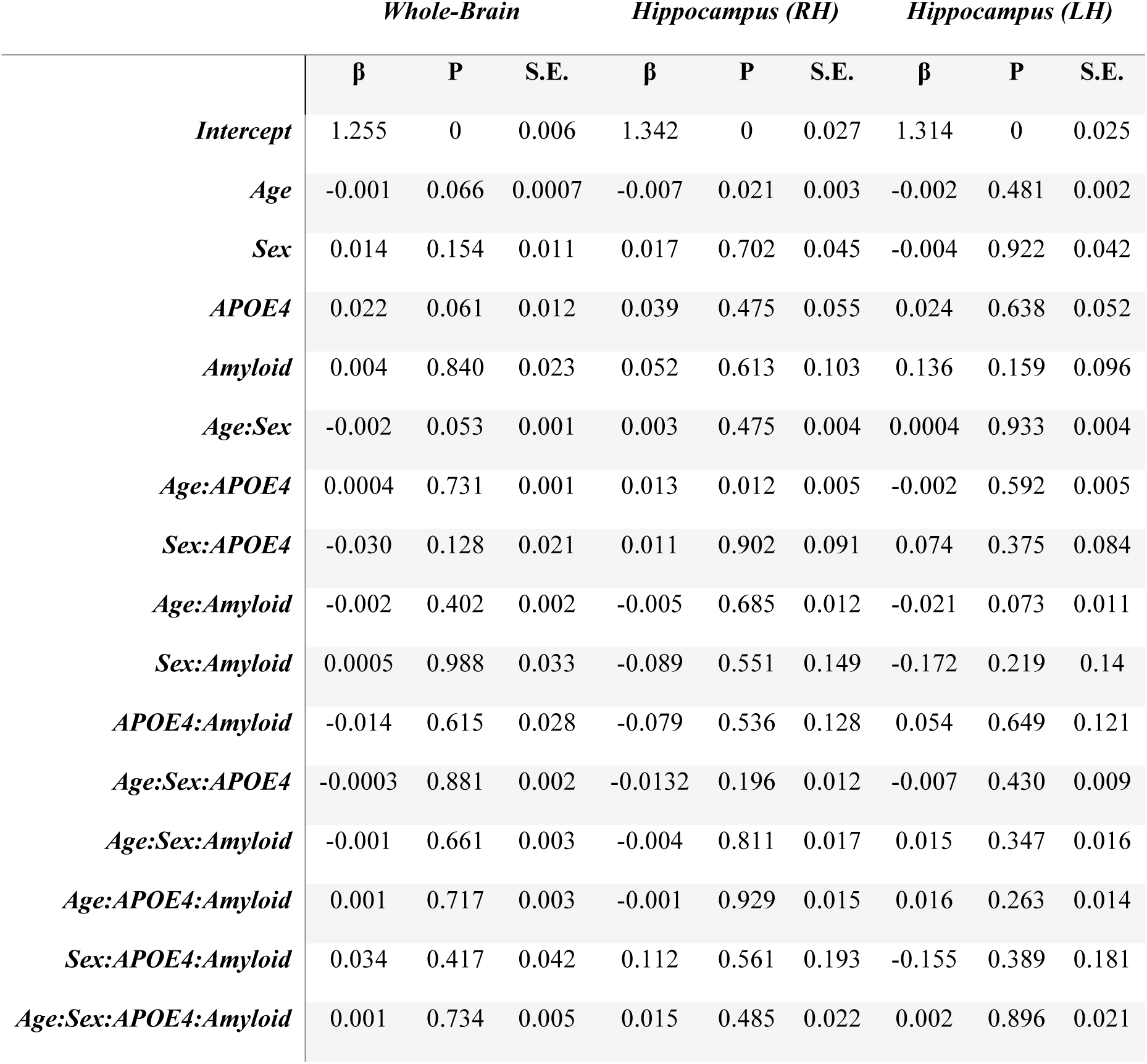
Regression model results examining EIB as a function of the interactions between age, APOE4, and amyloid (allowing for up to 4-way interactions between predictors), as well as two-way interactions of APOE4 – amyloid when controlling for other confounds.

